# Differentiated SH-SY5Y Cells Exhibit Neuronal Features but Lack Synaptic Maturity

**DOI:** 10.1101/2025.07.21.665912

**Authors:** Jana Leuenberger, Grischa Ott, Thomas Nevian, Benoît Zuber, Iman Rostami

## Abstract

A vital question in neuroscience is if and how efficiently cellular models may be differentiated into functional neuronal cells in culture. Despite the frequent use of the human neuroblastoma cell line SH-SY5Y, differentiation protocols vary extensively, with the most common being differentiation via the addition of retinoic acid and brain-derived neurotrophic factor. However, due to the lack of a reliable evaluation method, their adequacy as synaptic models remains unclear. Here, we investigate whether SH-SY5Y cells constitute a functional model for synaptic studies by phenotypically and ultra structurally analyzing synaptogenesis in SH-SY5Y cells subjected to different differentiation protocols. Electron microscopy (EM) techniques, including conventional EM, cryo-EM, and cryo-electron tomography, were systematically applied to characterize synaptogenesis in SH-SY5Y cells. Further characterization was performed through immunostaining and functional assays, such as live exocytosis assays and whole-cell patch-clamp electrophysiology. Despite exhibiting some presynaptic-like features, differentiated SH-SY5Y cells do not form morphologically or functionally complete synapses under the conditions tested. Immunostaining results were consistent with previous findings, showing synaptic markers. However, functional investigations did not detect synaptic activity. High-throughput EM analyses revealed an absence of synaptic structures in these cells. Additionally, an alternative differentiation approach incorporating additional neurotrophic factors promoted the formation of pre-synaptic-like compartments containing synapse-like synaptic vesicles (SVLVs). Though these SVLVs exhibited pleomorphic size distributions, differing from typical synaptic vesicles, and lacked connectors. These findings emphasize the need for cautious interpretation of results derived from SH-SY5Y cells when studying molecular synaptic architecture or neurodegenerative diseases.

## Introduction

There are numerous neuronal cell models, among the most used ones are human neuroblastoma cell lines, such as SH-SY5Y, mouse neuroblastoma cell lines such as NB41A, Neuro2a and dopamine-containing hybrids (MN9D), used in neuronal differentiation, neurotoxicity, neurodegenerative diseases and cancer ^1–7^. Other widely used cell lines include rat derived cells, such as catecholaminergic cells (PC12) ^8–10^, and human embryonic neuronal precursors (LUHMES) ^11^. Furthermore, induced pluripotent stem cell (iPSC)-derived neurons are increasingly regarded as an alternative model for studying synapses due to their potential to differentiate into diverse neuronal subtypes and form structures that more closely resemble native neurons ^12,13^. However, the production of iPSC-derived neurons is resource-intensive, requiring specialized expertise and infrastructure, expensive culture media, and long differentiation times. This and limited scalability make them less suitable for high-throughput applications compared to other models, like SH-SY5Y cells. As such, the choice of model often reflects a balance between physiological relevance and experimental feasibility. SH-SY5Y cells provide a faster and more cost-effective alternative for investigating neuronal pathways. While their structural and functional complexity does not fully match that of primary neurons or animal models, their use facilitates scalable and ethically sustainable experimentation. Although primary neurons remain the gold standard for synaptic studies, they are constrained by limited availability, variability, and ethical concerns, further underscoring the need for standardized, accessible cell models such as SH-SY5Y.

Synapse formation is a key determinant of neuronal maturation and function and a prerequisite for cellular models intended to investigate synaptic structure and plasticity ^14^. Synaptic model systems are vital to understanding signal transmission, circuit formation, and mechanisms underlying neurological disorders ^15^. SH-SY5Y cells, derived from human neuroblastoma, have been extensively used to model neuronal differentiation, function, and disease ^16–18^. They are known to express synaptic proteins, to exhibit neurite outgrowth, and evoked electrical activity or electrophysiological responses ^19–22^. However, whether these features translate into the formation of mature synaptic structures remains unclear.

High-resolution techniques such as transmission electron microscopy (TEM) and cryo-electron tomography (cryo-ET) offer direct visualization of synaptic ultrastructure and are essential for confirming synapse formation ^23,24^. Despite their potential, these methods remain underutilized in the characterization of SH-SY5Y-derived neuronal models.

This gap is particularly relevant considering the numerous differentiation protocols designed to enhance the neuronal properties of SH-SY5Y cells. Differentiation of SH-SY5Y cells typically requires specific cell culture medium supplementation. The most common supplement is all-trans retinoic acid (RA), a factor promoting early neuronal development through the activation of nuclear receptors ^25–28^. Supplementation with additional agents, such as phorbol esters and brain-derived neurotrophic factor (BDNF), enhance maturation and dopaminergic phenotype ^29,30^. Other supplements, such as dibutyryl-cAMP, promote neurite extension while cholesterol supports synaptic vesicle formation ^31–33^. Recent protocols combining RA, BDNF and B-27™ in neurobasal medium have also been reported to enhance neuronal differentiation of SH-SY5Y cells ^19^.

In this study, we evaluated the structure and functionality of SH-SY5Y cells, differentiated over 28 days, using live confocal microscopy, protein expression analysis, transmission electron microscopy (TEM), cryo-electron tomography (cryo-ET) and whole-cell patch clamp recordings. We report that while differentiation increased synaptic protein expression, no protocol, including RA, BDNF, cAMP, or cholesterol supplementation, led to mature synapse formation. This highlights the limitations of existing protocols to successfully differentiate SH-SY5Y cells into a functional synaptic model and underscores the need to explore alternatives.

## Methods

### Cell Culture

SH-SY5Y cells were maintained in Eagle’s minimum essential medium (EMEM, Gibco, Thermo Fisher Scientific), supplemented with 10% heat-inactivated fetal bovine serum (hiFBS, Gibco) and 1% penicillin-streptomycin (P/S, Gibco) in T-75 flasks, incubated at 37°C 5% CO_2_. The cells were passaged at a 1:10 ratio every week using 3mL 0.25% Trypsin-EDTA (1X) (Gibco, Thermo Fisher Scientific). Glass coverslips (BRAND, Sigma-Aldrich) and holey carbon-coated gold EM grids (300 mesh with Quantifoil R 2/1 or 200 mesh with lacey carbon film, EMS) were placed into dishes and were pre-coated with diluted (1:20 with ddH2o) 0.01% Poly-L-Lysin (PLL, Sigma-Aldrich)^16^. Cells were seeded at a density of 100,000 cells per well of a 6-well plate in EMEM supplemented with 10% hiFBS, 2mM Glutamine and 1% P/S. They were further differentiated using the media compositions adapted from Shipley et al. (2016). At day 3, the medium was exchanged to differentiation medium #1 (EMEM, 2.5% hiFBS, 2mM Glutamine, 1% P/S 10 µM RA (Sigma-Aldrich)). At day 5 Differentiation medium #2 (EMEM, 1% hiFBBS, 1% P/S, 2mM Glutamine and 10 µM RA) was introduced and at day 7 differentiation medium #3 (Neurobasal (Gibco, Thermo Fisher Scientific), 1X B-27 (Gibco, Thermo Fisher Scientific, 20mM KCl, 1% P/S, and 2mM Glutamax (Gibco, Thermo Fisher Scientific)) was used and maintained until day 14. On day 14, SH-SY5Y cells were further differentiated by adding 50 ng/mL BDNF (STEMCELL) or 50ng/mL BDNF, 10 ng/mL GDNF (STEMCELL), 10 ng/mL CNTF (STEMCELL) and 10 ng/mL IGF1 (STEMCELL) to medium #3. The medium was exchanged every three days until day 28.

### Immunofluorescence

Cells grown on coverslips were removed from the dish and were fixed with 3% paraformaldehyde and 0.5% glutaraldehyde (Agar Scientific, UK) for 1h at room temperature (RT) followed by washing in PBS three times. The cells were permeabilized with 0.3% Triton X-100 (Sigma-Aldrich) for 15 mins and washed again with PBS. 3% SureBlock (LubioScience) in PBS was used for unspecific blocking for 1h at RT. The cells were labelled with primary antibodies overnight and subsequently secondary antibodies (see Supplemental Table) in 3% blocking agent for 1h at RT, washing in-between 3x 5 min with PBS. The coverslips were dried overnight at 4°C. The coverslips were mounted on glass slides using Antifade mounting medium (ProLong Glas, Thermo Fisher Scientific) and were stored at 4°C until imaged.

### Immunoblotting

For immunoblots, SH-SY5Y cells were cultured as mentioned above on a 60mm dish at a seeding density of 250,000 per dish. The cells were washed twice with PBS before adding 400 µL RIPA buffer with 1:100 EDTA-free proteinase inhibitor (Sigma-Aldrich). They were lyzed for 30 min on ice on a shaker. They were then detached with a cell scraper and transferred into 1.5mL tubes. The lysate was then sonicated 3x 10 s at 50 % power using the SONOPLUS HD2070 homogenizer (Bandelin) and incubated on ice for further 10 min. The fully lyzed cells were centrifuged at 14.5 x g for 20 min and the supernatant containing the proteins of interest was collected. The protein content was determined using a bicichoninic acid assay (Sigma-Aldrich). The samples were prepared in Laemmli buffer, boiled0020for 5 min at 95°C, snap frozen, and stored at -80°C. The samples were equilibrated to 5 µg/well and loaded onto a 10% bisacrylamide gel. The gel was run for 30 min at 20mA, followed by 90 min at 40 mA. The gel was then transferred onto a nitrocellulose membrane (Merck) for 70 min at a constant 100 V. The membrane was washed with phosphate buffered saline with 1% Tween (PBST) for 10 min and then incubated with Intercept (PBS) Blocking Buffer (LI-COR Biosciences) in PBST for 1h. The primary antibodies (see Supplemental Table 1) were added in the blocking buffer overnight, washed 3x with PBST and then further incubated with HRP-conjugated secondary antibodies (See Supplemental Table 1) for 1 h. The washed membranes were incubated with ECL Reagent (Thermo Fisher Scientific) detected using the Fusion Fx system (Vilber Lourmat). Western blot images were quantified using GelAnalyzer and compiled in Excel (Microsoft Corporation).

### Vesicle recycling assay

The vesicle recycling assay was adapted from Gaffield & Betz (2007) and Iwabuchi et al. (2014) using the FM dye AM4-64 (Biotium, VWR International)^34,35^. To enable gentle medium exchange during imaging, a custom-made perfusion system using 20-gauge needles adapted to a µ-dish. The perfusion system was installed in the LSM 880 confocal microscope (Zeiss). Cells were washed in Tyrode’s solution (124 mM NaCl, 5mM KCl, 2 mM CaCl_2_, 1 mM MgCl_2_, 30 mM glucose, 25mM HEPES; 310 mOsm/l and pH 7.4) and then incubated for 2 min in Tyrode’s solution containing 10 µM AM4-64 dye. The cells were stimulated for 2 min with a depolarizing high-potassium solution (Tyrode’s containing 70 mM KCl, 59 mM NaCl and 10 µM AM4-64). This was followed by a 10 min incubation in standard Tyrode’s solution containing 10 µM AM4-64 to allow slow endocytosis staining (Gaffield & Betz, 2007). Non-internalized dye was quenched for 5 min with 0.5 mM SCAS in low-calcium and high-magnesium Tyrode’s solution (0.2 mM CaCl_2_ and 5 mM MgCl_2_), which halts spontaneous re-exocytosis of the endocytosed vesicles. A z-stack was acquired at the region of interest. Then the cells were stimulated with high-potassium Tyrode’s solution and a micrograph was acquired every 2 s for 3 min using a large pinhole (5.42 AU), thus broadening fluorescence detection in the z-plane (depth of focus 0.99 μm) using the laser at 488 nm and 1.8% strength. This allowed us to minimize laser intensity and reduce photobleaching. A z-stack of the region of interest was acquired after stimulation.

### Whole-cell patch-clamp electrophysiology

SH-SY5Y cells were cultured on 12 mm round glass coverslips (Fisher Scientific) and were transferred to a recovery chamber filled with oxygenated and carbonated artificial cerebrospinal fluid (aCSF) containing 125mM NaCl, 2.5mM KCl, 25mM NaHCO_3_, 1.25mM NaH_2_PO_4_, 1mM MgCl_2_, 2mM CaCl_2_, and 25mM glucose, kept at RT. Whole-cell patch-clamp recordings were performed using heat-pulled borosilicate glass pipettes with a tip resistance of 4–9 MΩ. The cells were visualized through an infrared charge-coupled detector camera mounted on a Leica DM LFSA microscope (Thorlabs). Cells were subjected to 500 ms square current injections from -60 to 300pA, in 20pA increments. Custom-made Igor Pro procedures (WaveMetrics) were used for all data acquisition and analyses ^36^.

### Electron microscopy of resin-embedded samples

The cells were fixed in 0.15 M HEPES containing 2.5% glutaraldehyde (Fluka) (670 mOsm, pH 7.35) at 4°C for a minimum of 24 h. They were then washed with 0.15 M HEPES three times for 5 min, post-fixed with 1% OsO4 (EMS) in 0.1 M Na-cacodylate-buffer (Merck) at 4°C for 1 h. Then, the cells were washed in 0.1 M Na-cacodylate-buffer three times for 5 min and dehydrated through a graded ethanol series (70, 80, 96 and 100% ethanol), each for 15 min at RT. Subsequently, the samples were infiltrated with a 1:1 mixture of ethanol and Epon 812 (Fluka) overnight at RT. They were then embedded in pure Epon 812 and left to harden at 60°C for 5 days. The resin blocks were removed from the dishes and sectioned with a UC6 ultramicrotome (Leica Microsystems), starting with semi-thin sections of 1 µm thickness which were stained with a 0.5% toluidine blue O solution (Merck). Ultrathin sections of 70nm thickness were cut with the ultra-diamond knife 45° (DiATOME). The sections were mounted on uncoated 200 mesh copper grids (G2200C, Plano GmBH), stained with uranyLess (EMS) and 3% lead citrate (Leica) using the EM STAIN device (Leica Microsystems, Vienna, Austria). The sections were imaged using a Tecnai Spirit transmission electron microcope (Thermo Fisher Scientific) equipped with a Veleta CCD camera (EMSIS) at an accelerating voltage of 80kV.

### Cryo-electron microscopy and tomography

SH-SY5Y cells were cultured on poly-L-lysin coated holey carbon film EM grids: R 2/1 – 300 mesh (Quantifoil) or lacey carbon film – 200 mesh (EMS). After transferring the grid to a custom-made plunge freezer, 4 µL of a solution containing 10 nm colloidal gold beads as fiducial markers was added. The grids were manually blotted using 12/N filter paper (Munktell) and vitrified by rapid plunging in liquid ethane and stored in liquid nitrogen. Cryo-EM micrographs were acquired using the FEI Tecnai F20 (200kV), with an FEI Falcon 2 direct detector. Tomograms were acquired on a Titan Krios G4 transmission electron microscope (Thermo Fisher Scientific) operating at 300 kV and 26,000x magnification. Tilt series were recorded from ±60° with 2° or 3° increments using the tomo5 software. The total electron dose per tilt series was adjusted to 1-1.55 e^-^/Å^2^, and images were collected at a target defocus of −10 µm. Data acquisition was performed using a Falcon 4i direct electron detector in combination with a Selectris energy filter. The tomograms were reconstructed using IMOD ^37^.

## Results

### SH-SY5Y differentiation pattern and synaptic proteins upon differentiation

The formation of synaptic connections is an essential prerequisite for neurons to reach functional maturation ^38^. Thus, we analyzed synaptic protein expression over time after initiating differentiation. SH-SY5Y cells were cultured and differentiated following established protocols, using differentiation media supplemented with cAMP, RA and BDNF for up to 28 days post-differentiation (DPD) ^3^. The cells were fixed and fluorescently labelled with synaptophysin, PSD95, β-tubulin III and DAPI and further processed for data acquisition at DPD 7, 14, 21 and 28. The cells showed progressive neurite outgrowth over time (Figure 1A1-4, S2-5), consistent with previous findings ^39^. β-tubulin III, a neurite marker associated with mature neurons, displayed a homogenous distribution throughout all neurites, reflecting the consistent development of neurite outgrowth and arborization over time (Figure 1A1-4, Figure S1-5, Figure S5) ^40–42^. Synaptophysin, an integral membrane protein of synaptic vesicles (SVs) involved in vesicle trafficking and endocytosis, is widely used as a marker for presynaptic terminals ^43^. It is endogenously expressed in SH-SY5Y cells and was observed to increase over time, as shown both by immunofluorescence microscopy (Figure 1A1-4) and western blot analysis (Figure 1B1). Quantitative western blot analysis (normalized to undifferentiated SH-SY5Y cells D0) revealed a 9.4 ± 2.5-fold increase in synaptophysin expression by DPD 28 (Figure 1B2), consistent with previous observations ^23^. Synapsin I, one of the most abundant presynaptic proteins in the brain, was detected at high levels, with a quantative increase of up to 17.3 ± 0.5-fold relative to expression at D0 (Figure 1B2) ^44^. The steep increase in this key presynaptic protein underscores the shift toward a more neuron-like protein expression pattern. In contrast, PSD95, a postsynaptic marker localized to the post synaptic density and known to regulate excitatory synapse maturation, was observed in comparatively low levels (Figure 1A1-4, B1) ^45^. Quantitative analysis showed a modest 1.4 ± 0.5-fold increase over the course of 28 days (Figure 1B2), suggesting a less pronounced development of potential postsynaptic structures.

**Figure 1.**
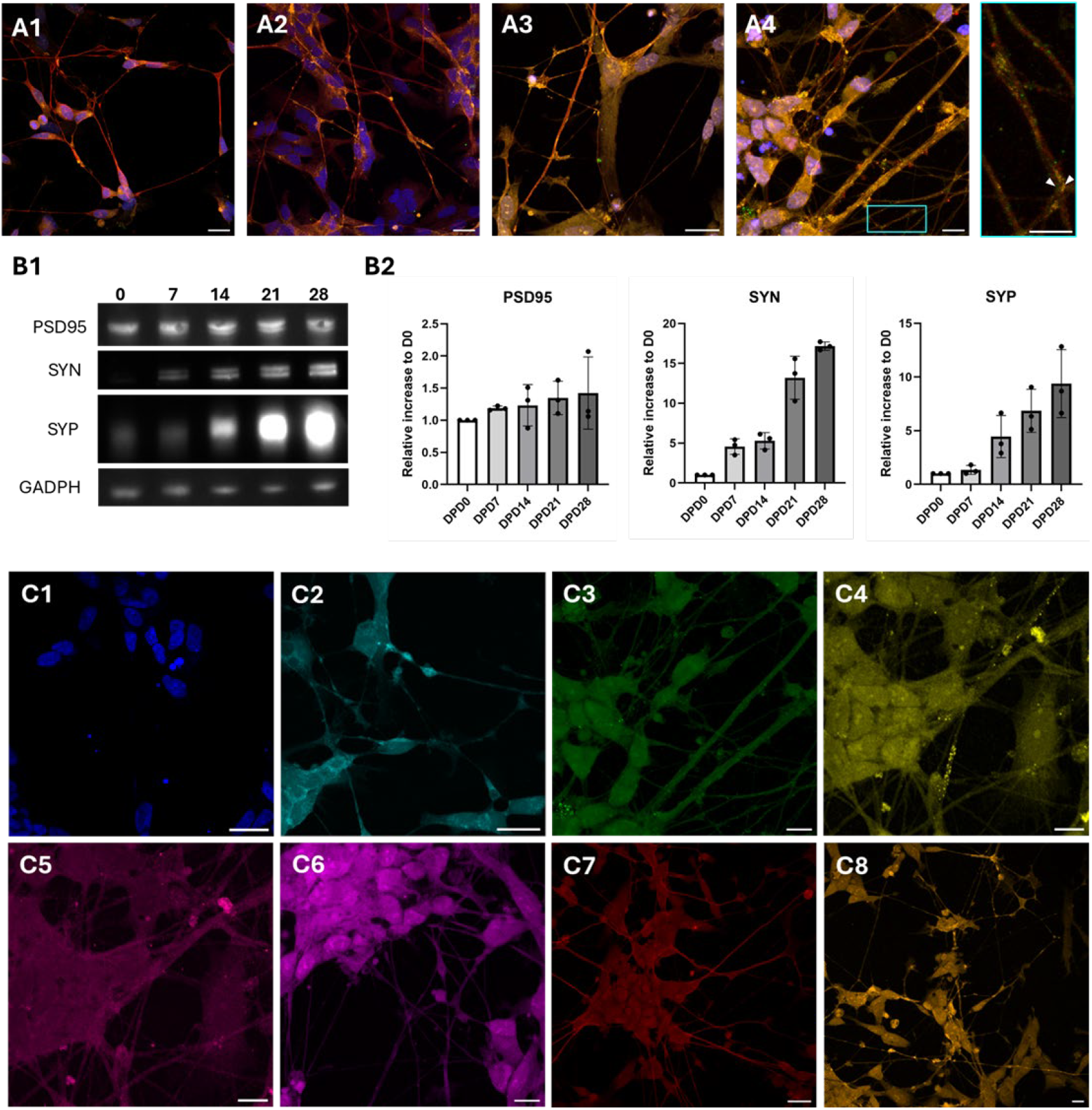
Relative increase in synaptic proteins. A: Confocal images of SH-SY5Y cells over a 28-day differentiation period at timepoints DPD 7 (A1), DPD 14 (A2), DPD 21 (A3) and DPD 28 (A4) showing an increase of the presynaptic marker synaptophysin (orange), the postsynaptic marker PSD95 (green), the neurite marker β-tubulin III (red), and DAPI (blue). Zoomed in square in cyan showing synaptic puncta (white arrows). Scale bars: 20µm and 5 µm (zoom-ins). B1: Western blot of the synaptic proteins PSD95, synapsin I and synaptophysin, as well as GAPDH (loading control) in undifferentiated cells (DPD 0) and over the course of differentiation (DPD 7-28). B2: Semi-quantitative analysis of the relative protein expression of PSD95, synapsin I (SYN) and synaptophysin (SYP) normalized by GADPH and expression at DPD0. C: Antibody stains of differentiated SH-SY5Y cells at DPD 28. C1: DAPI, C2: Synapsin I, C3: PSD95, C4: vGLuT1, C5: GluA2, C6: Rab3a, C7: β-tubulin III, C8: Synaptophysin. Scale bars: 20µm.^52^

A characteristic feature of mature neurons is the clustering of synaptic proteins along neurites and at sites of synaptic contacts. When fluorescently labelling these clusters, they appear as discrete puncta. Labeling of both pre- and postsynaptic proteins typically shows puncta that are closely apposed or overlapping, which can serve as an indicator of synapse formation and density ^46^. Synaptophysin displayed a mild punctate pattern, while PSD95 formed fewer visible puncta (Figure 1A1-4; merged in Figure S2-5, Figure 1C3, C8). However, little to no spatial proximity between pre- and postsynaptic puncta was observed (Figure 1A4, zoom-in). PSD95 puncta were sparse at both DPD 21 and 28 (Figure 1A3, A4, unmerged in Figure S4, S5), and their lack of overlap with synaptophysin puncta suggests a low degree of synapse formation, particulary when compared to mature neuronal cultures, such as the ones found in hippocampal cultures ^47^.

Beyond synaptophysin, PSD95 and β-tubulin III, we examined the localization of additional key synaptic proteins to gain a more comprehensive overview of SH-SY5Y cell neuron-like characteristics. These include the presynaptic markers synapsin I, vGLuT1 and Rab3a, as well as the post-synaptic GluA2. Synapsin I exhibited a homogenous distribution throughout the cytoplasm with slightly elevated signal intensities at varicosities along neurites (Figure 1C2). However, no clear clustering into presynaptic puncta was observed. vGLuT1, a presynaptic vesicular transporter, showed localized puncta along neurites, suggesting glutamatergic properties in differentiated SH-SY5Y cells (Figure 1C4) ^48–50^. Rab3a, a small GTPase localized at presynaptic terminals, showed a diffuse distribution throughout the cytoplasm (Figure 1C6) ^51^. Similar to PSD95, GluA2, a subunit of AMPA receptors, primarily localized at postsynaptic sites, did not display punctae supporting the notion of limited postsynaptic formation (Figure 1C5) ^52^.

### SH-SY5Y cells Differentiated by RA and BDNF Do Not Form bona fide Synapses

To investigate the ultrastructural features of differentiated SH-SY5Y cells, we analyzed neurites and their terminal specializations using both conventional TEM and cryo-ET. Cells were either seeded in 6-well plates for conventional EM or cultured on EM grids cryo-ET. We focused on neurites and their and their swellings, putative axonal boutons, as potential sites of synaptic specialization (Figure 2A). We analyzed over 1000 conventional EM micrographs, more than 500 cryo-EM images and over 500 cryo-ET reconstructions. These data revealed extensive neurite outgrowth and frequent cellular connections (Figure 2C). SH-SY5Y axons typically exhibited a uniform diameter with densely packed microtubules and limited branching compared to dendrites (Figure 2C). Interspersed among the microtubules were 10nm-wide intermediate filaments (Figure 3B). Long tubular mitochondria ranging from tenths of micrometers to several micrometers were abundant within axonal neurites (Figure 2C). Actin-rich membrane protrusions lacking microtubules were identified at neurite tips and branching points, indicative of filopodia and growth cones (Figure 3D). These findings are consistent with early axon branching previously described in primary hippocampal neurons ^53,54^. Dendrites, identified by their larger diameter, the presence of spines, polyribosome clusters, and a less dense cytoskeletal organization, were less frequently observed^53^.

**Figure 2.**
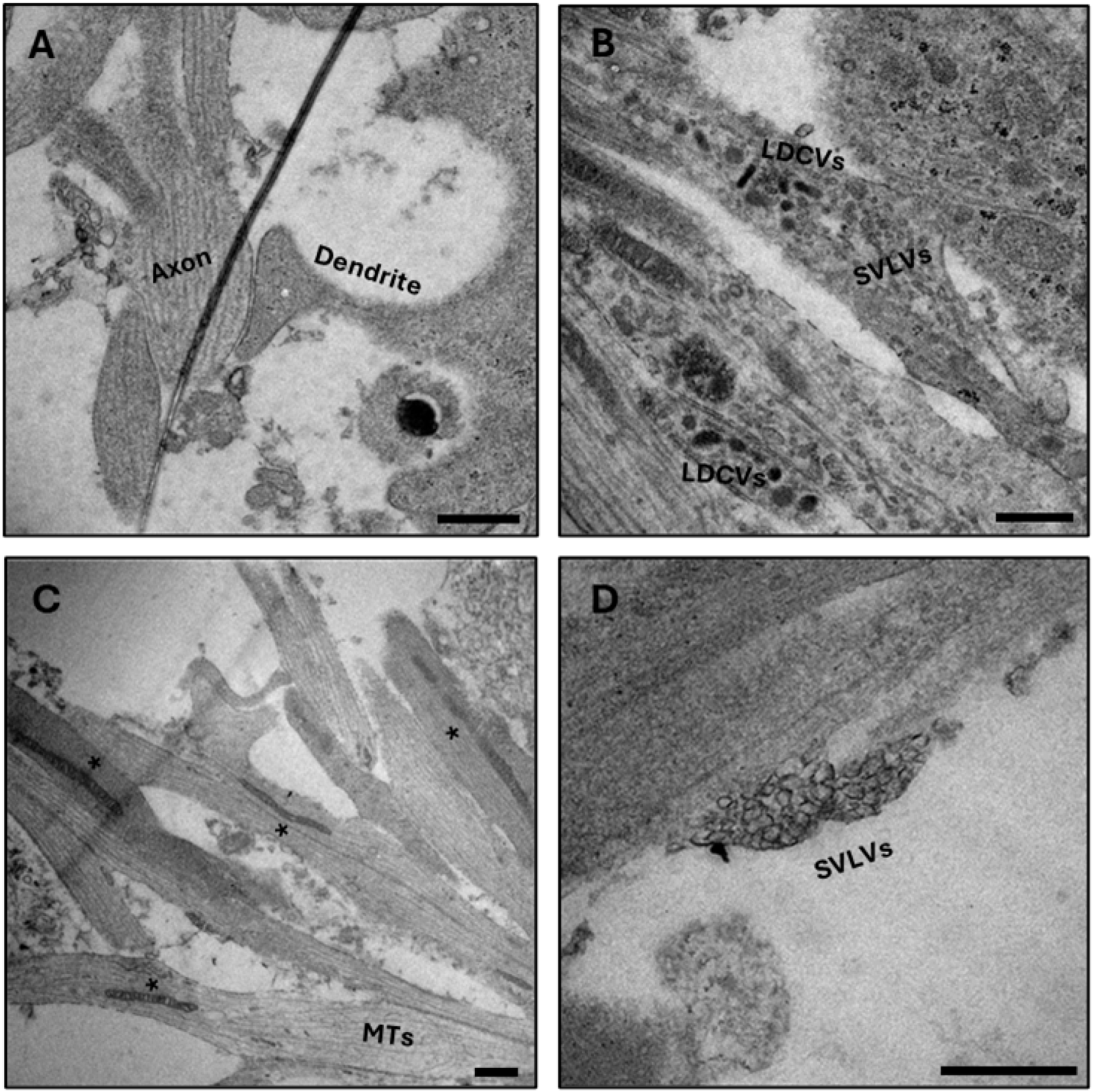
SH-SY5Y cell neuronal features in conventional electron microscopy. A: Synapse-like connection lacking SVs and active zone. B: Neurite estensions containing large dense-core vesicles (LDCVs). C: Long tubulular mitochondria (*) in microtubule (MT) containing axon bundles. D: Peripheral vesicle clusters with vesicles of similar size to SVs located along neurites. Scale bars: 200 nm.

**Figure 3.**
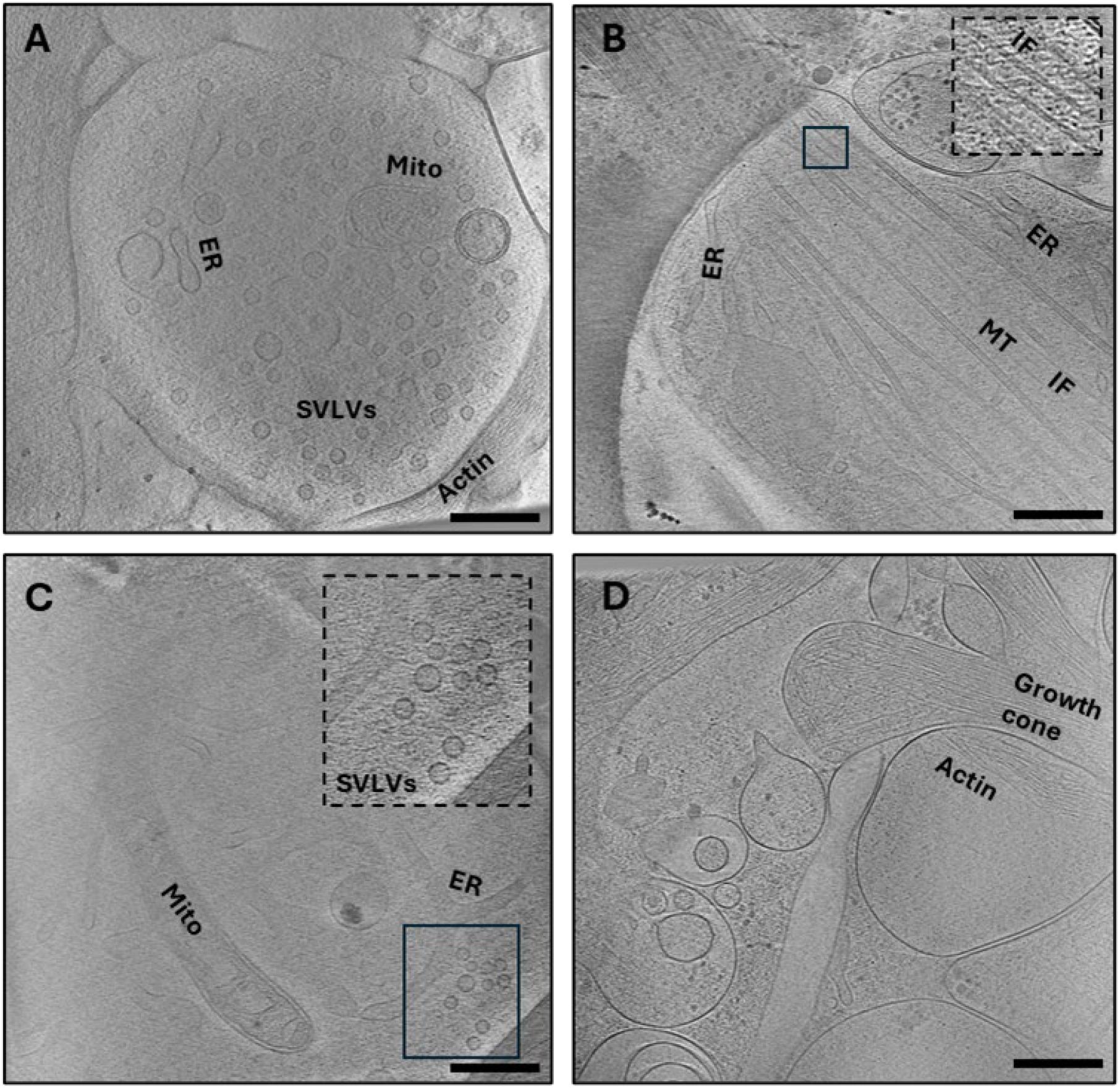
Cryo-electron tomogram of neuronal structures in SH-SY5Y cells. A: Pre-synaptic-like bouton containing SVLVs, mitochondria and endoplasmic reticulum (ER). B: Axonal neurite with microtubules, tubular ER and intermediate filaments (zoom-in). C: Small cluster of SVLVs (zoom-in), mitochondria and ER. D: growth cone with actin-filled filopodia of a newly forming neurite. Abbreviations: Mito: Mitochondria; ER: endoplasmic reticulum, SVLV: synapse-like synaptic vesicle, IF: Intermediate Filament; MT: Microtubule. Scale bars: 100 nm.

During differentiation, we observed fine cellular protrusions between SH-SY5Y cells, particularly at earlier points (DPD 14), that resemble tunneling nanotubes (TNTs) (Figure S11)^18^. These structures were less prominent at later stages (DPD 28), when neurite-like processes dominated morphology. Although these protrusions superficially resemble structures implicated in intercellular interactions, such as thin membrane bridges, we did not assess cytoskeletal composition, membrane continuity, or intercellular transport, and therefore refrain from making further interpretations regarding their identity or function ^55,56^. Nevertheless, their presence in early-stage cultures may point to dynamic morphological features of SH-SY5Y cells that merit further investigation.

In primary neurons and synaptosomes, mature synapses are structurally characterized by a presynaptic bouton containing clusters of small, clear SVs, often arranged in distinct pools and interconnected by filamentous connectors. These boutons feature a defined active zone aligned with the synaptic cleft and a postsynaptic compartment, typically a dendritic spine ^57,58^. In differentiated SH-SY5Y cells, presynaptic-like structures were observed at axonal boutons containing SVLVs. These vesicles had a mean diameter of approximately 35 nm, which is slightly smaller than typical SV (∼45 nm) (Figure 2D; Figure 3A, C). These vesicles appeared to be devoid of any connectors, despite the presence of small clusters (Figure 3C, zoom-in), which would generally be found in SV pools. This implies the absence of a higher structural synapse organization (Figure 2D; Figure 3A,C) Additionally, the presynapse-like bouton lacked the characteristic active zone and a postsynaptic counterpart (Figure 3A, C). Similarily, varicosities along axons containing SVLVs and mitochondria are abundantly present. However, at these varicosities also no typical SV architecture, i.e. organized pools, connectors, tethers, and AZ were observed (Figure 2D). The SVLVs were of pleomorphic shape and size, which indicates immature presynaptic compartment formation. We further observed multivesicular bodies (MVBs), which may be easily mistaken for presynaptic terminals due to their small vesicles enclosed in a large membrane, but are distinguishable by the protein decoration on the vesicles (Figure S7). Large dense core vesicles (LDCV) presenting as 100 to 300nm dark vesicles were also seen in neurites as well as in close proximity to SVLVs (Figure 2B). As cholesterol has been implied to aid in SV formation, we further investigated SH-SY5Y cells differentiated by this approach using high trouphput screening of EM sections. Results showed no amelioration in synaptic ultrastructure and was thus not further evaluated (Figure S6).

### Adding additional neurotrophic factors did not further synapse maturation

We attempted to trigger a more mature neuronal state by supplementing the culture with additional neurotrophic factors (NTFs), namely glial-derived neurotrophic factor (GDNF), ciliary neurotrophic factor (CNTF) and insulin-like growth factor (IGF-1), which were subsequently added after DPD14 ^59–61^. These SH-SY5Y cultures were subjected to protein profile analyses by fluorescent staining and western blotting, ultrastructural investigations, as well as functional assays. Comparative western blot analysis of SH-SY5Y cells differentiated with RA and BDNF alone or additional NTFs showed a slight increases of vGLuT1 and SYP from DPD 21 to DPD 28 in both conditions, but no difference between the two groups (Figure 4C). Morphological analysis by immunofluorescence (Figure 4A, unmerged in Figure S10) revealed that SH-SY5Y cells treated with additional NTFs tended to exhibit more separated neurites in contrast to the prominent axon bundling observed with BDNF and RA alone (Figure 1A1-4). EM analysis showed that in spite of addional NTFs, presynaptic-like structures lacked a postsynaptic partner, AZ and vesicle connectors (Figure 4B). In conclusion, we observed no significant changes following supplementation with additional NTFs, indicating that they do not further promote neuronal maturation in SH-SY5Y cells.

**Figure 4.**
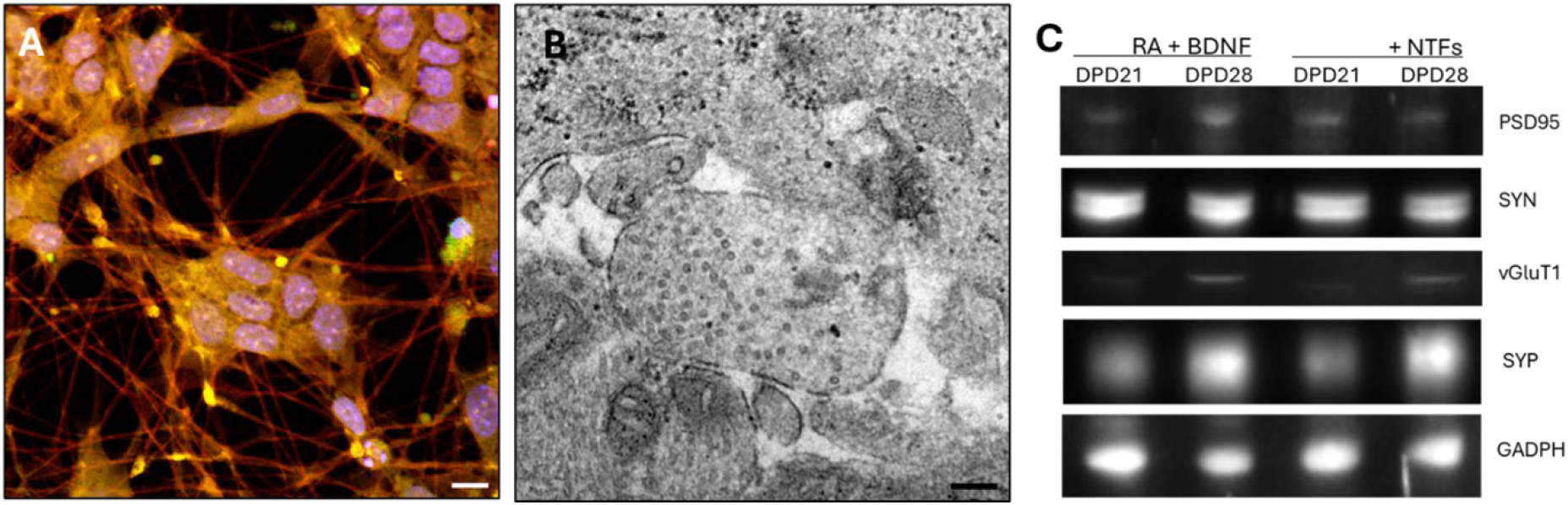
Analysis of differentiation by RA + BDNF with additional NTFs. A: Confocal image of NTF-differentiated cells stained with Synaptophysin (orange), PSD95 (green), β-tubulin III (red) and DAPI (blue). Immunofluorescence stain shows no clear synaptic puncta. Scale bar: 10 µm. B: Conventional EM micrographs of a presynapse-like bouton containing SVLVs. Scale bar: 200 nm. C: Western blot analysis of synaptic protein expression levels in RA+BDNF-differentiated compared to NTF-differentiated SH-SY5Y cells at DPD 21 and 28 of synaptophysin (SYP), synapsin-I (SYN), PSD95, vGluT1 and GADPH (loading control) showed no significant difference in protein expression level.

### Synaptic activity cannot be achieved by common differentiation methods

To evaluate the functional maturation of SH-SY5Y cells, we performed whole-cell patch clamp recordings in both current clamp and voltage clamp configurations across four experimental conditions, DPD 21 and DPD 28, with and without neurotrophic factor supplementation. Current-clamp recordings demonstrated single action potentials in response to stepwise depolarizing current injection (from -60 to +160 pA in 20pA steps) (Figure 5). Voltage-clamp recordings, holding the cell at -60mV, showed no spontaneous postsynaptic potentials, indicating a lack of detectable spontaneous synaptic activity (Figure 5). The absence of depolarizing or hyperpolarizing events suggests that differentiated SH-SY5Y cells do not establish functional synaptic connectivity under the applied differentiation protocol. Notably, even prolonged differentiation and trophic support failed to induce functional excitability or voltage-gated conductances typical of mature neurons. These results indicate that despite extended culture and supplementation, differentiated SH-SY5Y cells do not attain electrophysiological properties characteristic of mature, excitable neurons.

**Figure 5.**
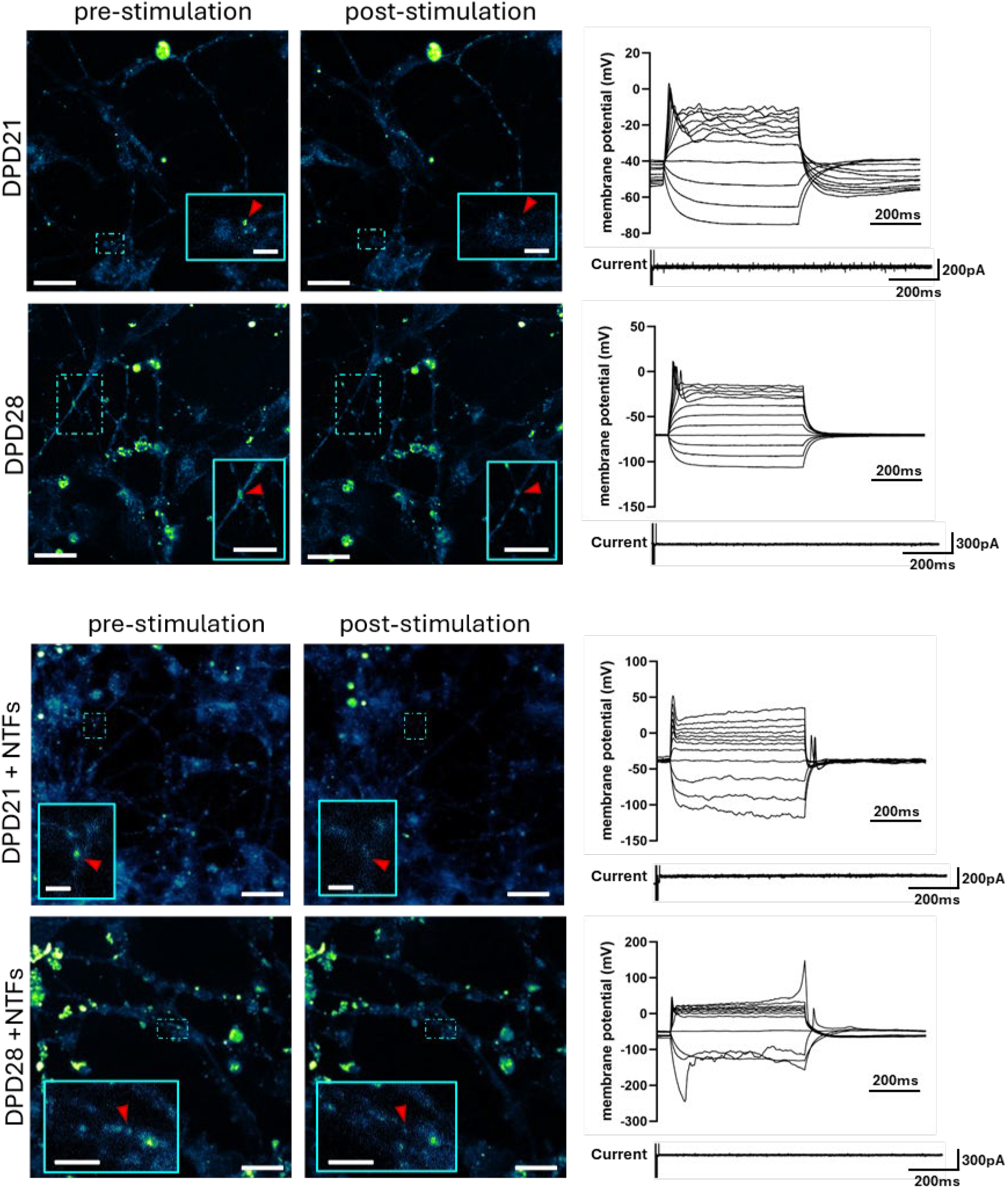
Functional live-stain assay with AM4-64 dye and electrophysiological traces. Fluorescence images of SH-SY5Y cells differentiated with either BDNF + RA at DPD 21 and 28 and with additional NTFs at DPD 21 and 28 stained with AM6-64 dye. Z-projected images of ROI with small zoom-ins on puncta pre-stimulation (leftside) and z-projected images with zoom-ins on puncta acquired post-stimulation with KCl after 120s (rightside). Zoom-ins show the movement of labelled vesicles (red arrow). Scale bars: 10 µm and 2 µm (zoom-ins). Respective current-clamp mode recordings (right side on the top) to the corresponding immunofluorescence images demonstrated singular action potentials in response to stepwise depolarizing currents. y-axis shows membrane potential (mV); x-axis shows time (ms) Voltage-clamp recordings (right side on the bottom) showed no pontaneous post-synaptic potentials (sPSPs), suggesting no signals received from neighboring cells. x-axis shows time (ms); y-axis shows spontaneous currents (pA).

To further assess stimulus-evoked exocytosis, we performed live-cell imaging using the fluorescent dye AM4-64, which binds reversibly to the plasma membranes and is internalized by endocytic events. subsequent quenching of non-internalized dye enables visualization of endocytosed vesicles. In neurons, recently endocytosed vesicles have a higher probability to undergo evoked exocytosis^62^. SH-SY5Y cells were assessed at DPD 21 and DPD 28. They were imaged after dye loading and quenching at 2-second intervals during high-potassium stimulation. While some fluoresenct puncta were observed in neurites at DPD 21 and increased in number at DPD 28, their apparent disappearence following stimulation was not due to vesicle exocytosis (Figure 5). Instead, time-lapse imaging revealed that vesicles remained fluorescent but moved along neurites and frequently shifted in and out of the imaging plane, which can mimick fluorescence loss (Figure S8, Figure S16-19).

## Discussion

In this study, we evaluated the neuronal properties of SH-SY5Y cells differentiated using established protocols, with or without supplementation with additional neurotrophic factors over a 28-day culture period. Our aim was to assess their suitability as an in-vitro neuronal model so study synaptic properties and processes. While differentiated SH-SY5Y cells exhibit several neuronal features, such as expression of presynaptic proteins, they do not recapitulate the structural and functional aspects characteristic of mature synapses.

Our findings indicate that SH-SY5Y cells undergo morphological development reminiscent of primary hippocampal neurons, characterized by rapid neurite outgrowth and expression of β-tubulin III, a marker of mature neurons. At the ultrastructural level, axons and dendrites can be distinguished, and organelles relevant in neuro- and synaptogenesis, such as mitochondria and ER, are present.

We observed that presynaptic and postsynaptic markers are expressed under standard differentiation protocols incorporating RA, BDNF, cAMP and B27. Key presynaptic proteins, including synapsin I and synaptophysin, and the postsynaptic scaffold protein PSD95 were abundantly detected. Some clustering of presynaptic proteins was observed; however, no corresponding synaptic puncta were shown to overlap. The diffuse distribution of most of these synaptic proteins resembles their distribution in immature neurons, in which synaptic proteins are often synthesized but not fully trafficked to synaptic sites, resulting in diffuse rather than punctate localization ^63^.

The organization of cytoskeletal elements provides insights into SH-SY5Y cell neuronal maturity. Ultrastructural analysis confirmed the presence of intermediate filaments, which are essential for maintaining axonal integrity and function ^64^. SH-SY5Y cells also exhibited actin-rich growth cones and distinct axon- and dendritic-like structures. In mature neurons, such cytoskeletal specializations are essential for synaptic function and neurite stability^65^. Closer examination on the cytoskeleton in Cryo-EM revealed thin extensions containing straight actin bundles, consistent with tunneling nanotubes (TNTs), which transport cellular material between cells (Figure S11) ^18^. These extensions were more abundant at early differentiation time points and were gradually replaced by axon-like neurites composed of microtubules. In SH-SY5Y cells, TNTs have been shown to mediate the transfer of α-synuclein ^66^, a protein implicated in Parkinson’s disease ^67,68^.

Despite extensive neurite outgrowth, SH-SY5Y cells failed to form functional synaptic connections. Although SVLVs were present at neurite terminals, they did not assemble into organized pools typically associated with mature presynaptic boutons ^57,58,69–71^. LDCVs were also observed. Taking together with the diffuse non-punctate distribution of synaptic proteins, these findings suggest that SH-SY5Y cells are unable to initiate synaptogenesis.

To further promote synaptogenesis, we differentiated SH-SY5Y cells in the presence of additional neurotrophic growth factors (GDNF, CNTF, IGF1), which have been reported to support neuronal maturation ^13,72^. However, biochemical and structural analyses revealed no significant enhancement in synaptic protein expression or synaptic ultrastructure. Electrophysiological recordings revealed that SH-SY5Y cells, despite additional NTF supplementation, exhibited only single action potentials, reflecting limited excitability likely due to insufficient ion channel density or overall neuronal immaturity ^73–75^. Such isolated action potentials are typical of immature neurons, in which ion channel kinetics and synaptic inputs are not yet fully developed ^76^. The lack of spontaneous synaptic activity in voltage-clamp recordings further supports the conclusion that SH-SY5Y cells lack functional maturation.

To investigate potential endocytic and exocytic activity, we used dye AM4-64 (S. O. Rizzoli & Betz, 2004; D’Aloia et al., 2024). Fluorescent puncta, corresponding to recently endocytosed vesicles, were observed in neurites, but rather than undergoing exocytosis upon repeat stimulation, they exhibited directional movements along neurites ^24,78^. This behavior reflects intracellular transport dynamics, rather than exocytosis^79^. The lack of neuron-like vesicle recycling relates to the absence of proper synapses in this cell model. As AM4-64 labels all membrane-bound compartments resulting from endocytosis, including early and recycling endosomes, lysosomes, and transport vesicles, the observed motility could be attributed to trafficking of these organelles ^80,81^.

We hypothesize that the failure of SH-SY5Y to undergo synaptogenesis may stem from a lack of target recognition mediated by cell adhesion molecules such as cadherins and integrins. These molecules are essential for establishing contact between pre- and postsynaptic sites ^82^.

In conclusion, our study demonstrates that SH-SY5Y cells differentiated with RA and BDNF fail to form synapses. Although these cells display several neuronal characteristics, their lack of functional synaptic connections and their immature electrophysiological properties limit their utility as a model for synaptic studies.

## Acknowledgments

This work was supported by grants from the Swiss National Science Foundation (31003A_179520, 32NE30_185536, CRSII-222809 to BZ). Images were acquired with microscopes of Dubochet Center for Imaging Bern (DCI-Bern) and the Microscopy Imaging Center (MIC) of the University of Bern. We thank Marek Kaminek and David Kalbermatter for their strong support in the usage of the electron microscopes. We also want to thank Beat Haenni and Desirée Dürr for the preparation of the conventional EM samples. Further, we extend our gratitude to Prof. Dr. Nevian’s group, especially Benjamin Leonardon and Mathilde Amat di San Filippo at the Institute of Physiology of the University of Bern for providing the facilities and expertise to enable our electrophysiological experiments.

