## Supplemental table, Supplemental figures for "Differentiated SH-SY5Y Cells Exhibit Neuronal Features but Lack Synaptic Maturity"

### Supplemental Materials

| Type | Descriptor / Conjugate | Manufacturer | Product number | Host | Dilutions used for IF / WB |
|---|---|---|---|---|---|
| **Primary Antibodies** | Anti-Synaptophysin | Sigma-Aldrich | S5768 | Mouse monoclonal | 1:1000 / 1:5000 |
| | Anti-Synapsin I | Sigma Aldrich | S193 | Rabbit polyclonal | 1:1000/ 1:5000 |
| | Anti-Rab3a | Proteintech | 16865-1-AP | Rabbit polyclonal | 1:1000 / 1:5000 |
| | Anti-vGluT1 | Synaptic Systems | 125511 | Mouse monoconal | 1:1000/ 1:5000 |
| | Anti-PSD-95 | Abcam | ab238513 | Rabbit monoclonal | 1:1000 / 1:1000-1:5000 |
| | Anti-PSD-95 | Invitrogen | MA5-45141 | Mouse monoclonal | 1:1000 / - |
| | Anti-MAP2 | Invitrogen | PA1-10005 | Chicken polyclonal | 1:1000 / 1:5000 |
| | Anti-GluA2 | Sigma Aldrich | ZRB1008 | Rabbit monoclonal | 1:1000/ 1:5000 |
| | Anti-β-III-tubulin | Novus Biologicals | NB100-1612 | Chicken polyclonal | 1:1000 / - |
| | Loading control: Anti-β-tubulin / HRP | Invitrogen | MA5-16308-HRP | Mouse monoclonal | - / 1:5000 |
| | Loading control: Anti-GAPDH / HRP | Invitrogen | MA5-15738-HRP | Mouse monoclonal | - / 1:5000 |
| **Secondary Antibodies** | Anti-Rabbit / Alexa-488 | Invitrogen | A11004 | Goat polyclonal | 1:1000 / - |
| | Anti-Rabbit / Alexa-568 | Invitrogen | A11011 | Goat polyclonal | 1:1000 / - |
| | Anti-Mouse / Alexa-568 | Invitrogen | A11004 | Goat polyclonal | 1:1000 / - |
| | Anti-Chicken / Alexa-647 | Jackson ImmunoResearch | 703-605-155 | Donkey polyclonal | 1:1000 / - |
| | Anti-Rabbit / HRP | Invitrogen | 31466 | Goat polyclonal | - / 1:5000 |
| | Anti-Chicken / HRP | ThermoFisher | A16054 | Goat polyclonal | - / 1:5000 |
| **Dyes** | DAPI | Sigma-Aldrich | D9542 | - | 1 µg/ml |
| | AM4-64 | Biotium | 70025 | - | 10 µM |
| **Quencher** | SCAS (4-Sulfonato calix [8] arene, sodium salt) | Biotium | 70037 | - | 0.5 mM |

*Table 1: Primary and secondary antibodies and dyes used*

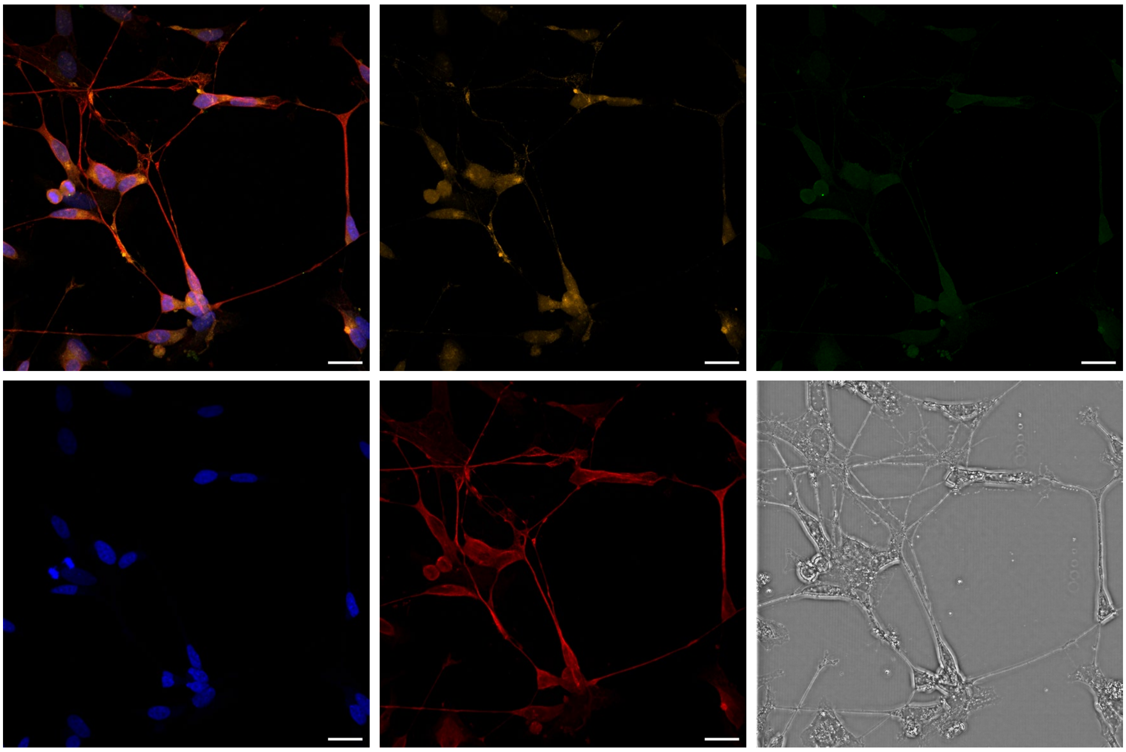

*Figure S1: Confocal images of SH-SY5Y cells corresponding to the merged image of DPD 7 in figure 1. Synaptophysin (orange), PSD95 (green), DAPI (blue), β-tubulin III (red) and brightfield (grey). Scale bars: 20 µm. Sparse staining of synaptophysin and PSD95 shows beginning differentiation towards a neuronal phenotype. Neurites start to elongate and connect to neighboring cells*

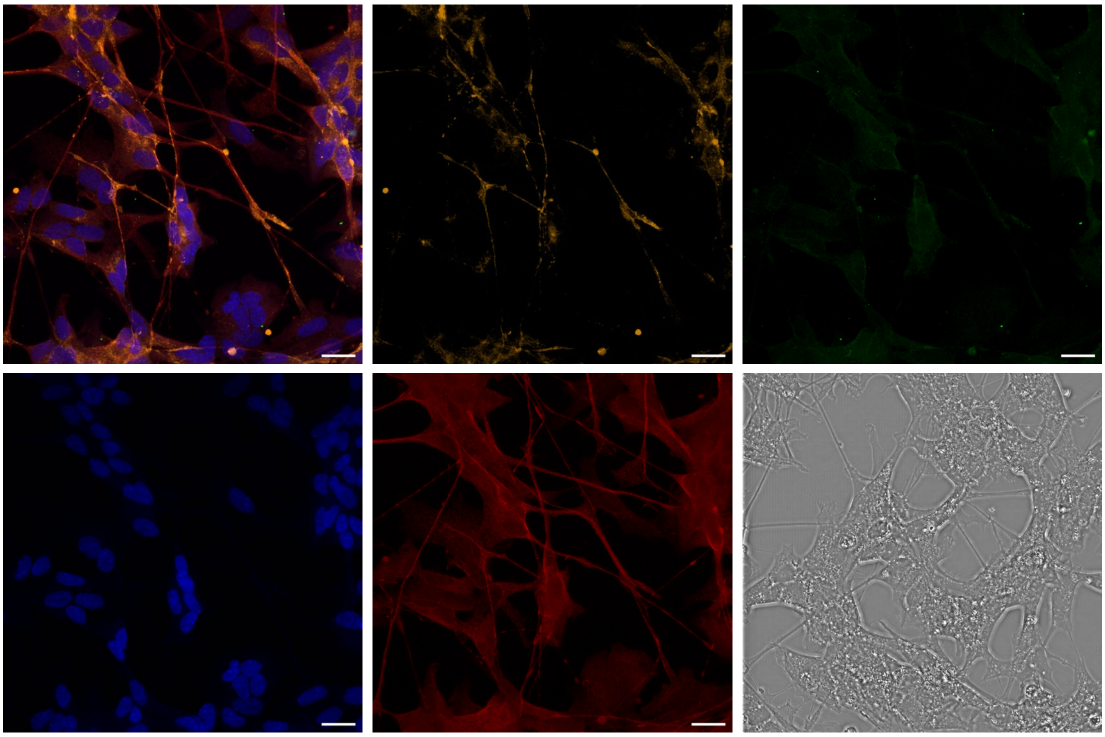

*Figure S2: Confocal images of SH-SY5Y cells corresponding to the merged image of DPD 14 in figure 1. Synaptophysin (orange), PSD95 (green), DAPI (blue), β-tubulin III (red) and brightfield (grey). Synaptophysin is seen in a more localized synaptic puncta manner, while PSD95 does not show a strong increase compared to DPD 7. Scale bars: 20 µm*

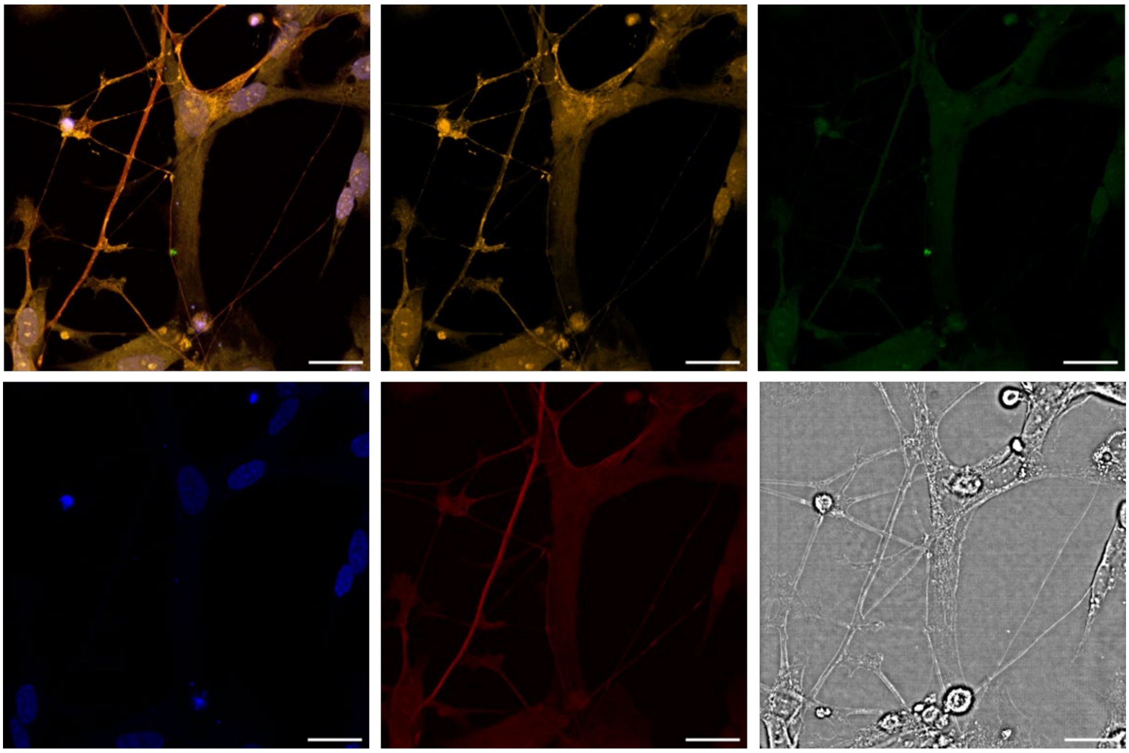

*Figure S3: Confocal images of SH-SY5Y cells corresponding to the merged image of DPD 21 in figure 1. Synaptophysin (orange), PSD95 (green), DAPI (blue), β-tubulin III (red) and brightfield (grey). Synaptophysin stain increases upon longer differentiation, while PSD95 does not increase. β-tubulin III becomes more localized during maturation. Scale bars: 20 µm*

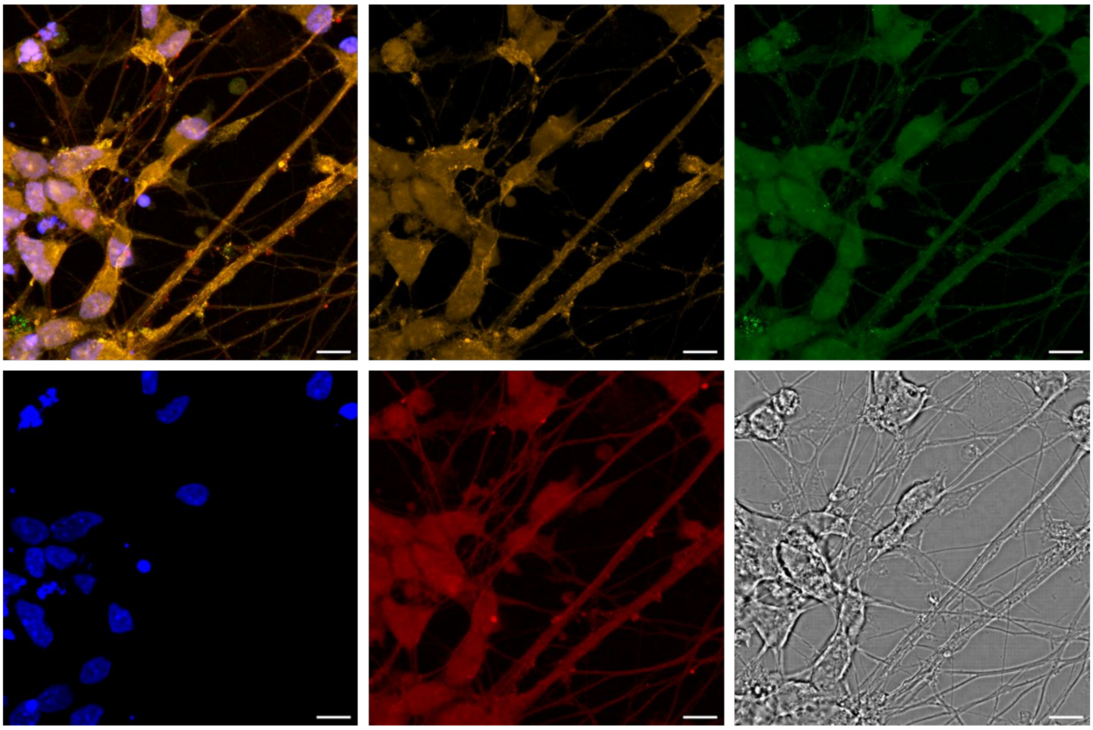

*Figure S4: Confocal images of SH-SY5Y cells corresponding to the merged image of DPD 28 in figure 1. Synaptophysin (orange), PSD95 (green), DAPI (blue), β-tubulin III (red) and brightfield (grey). Presynaptic puncta become more abundant and slight post-synaptic puncta are present indicating further maturation. Axon bundles form and present in a more branching manner and appear more pronounced. Scale bars: 20um*

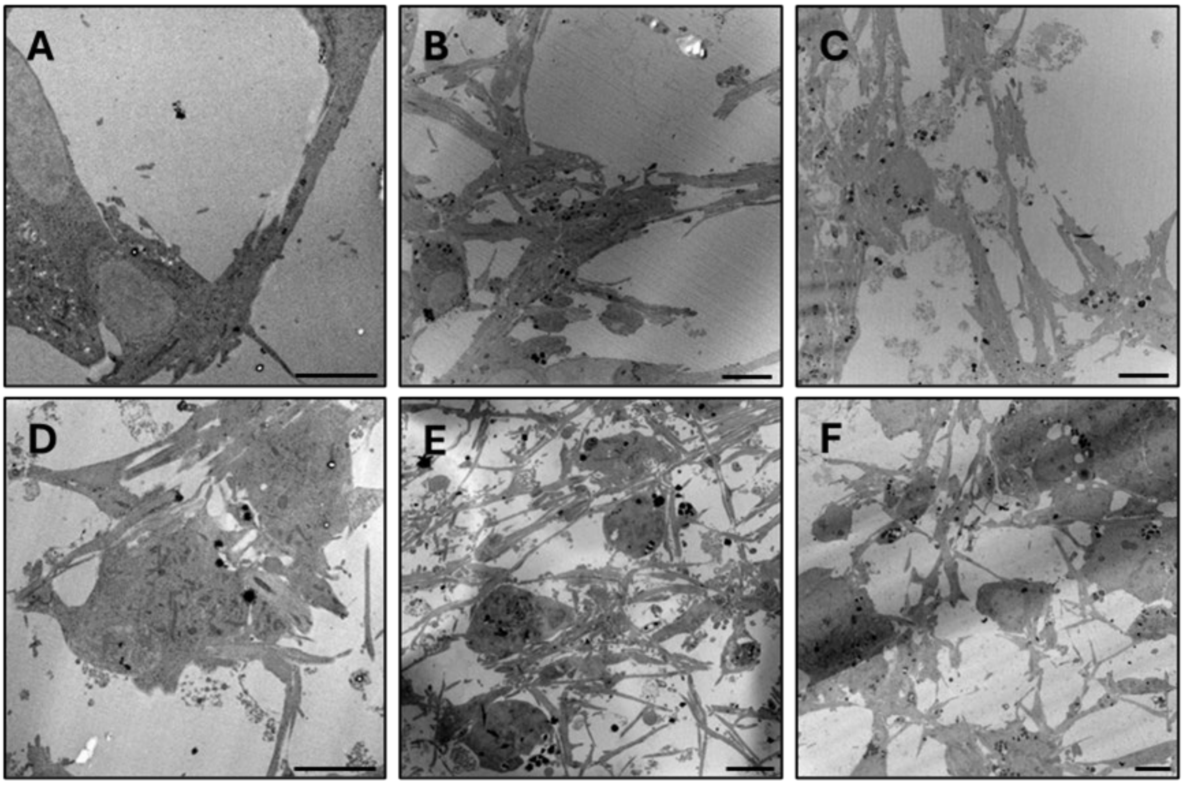


*Figure S5: Electron micrographs of SH-SY5Y cells at A: DPD 7, D: DPD 14, B: DPD 21, and C: DPD 28 differentiated using RA + BDNF and E: DPD 21 and F: DPD 28 differentiated using NTFs. Neurites increase in length and number over differentiation period gradually with a slightly more bundled phenotype in cultures differentiated with RA + BDNF in comparison to the more separated neurite network observed upon addition of the NTFs CNTF, GDNF and IGF1. Scale bars: 5 µm*

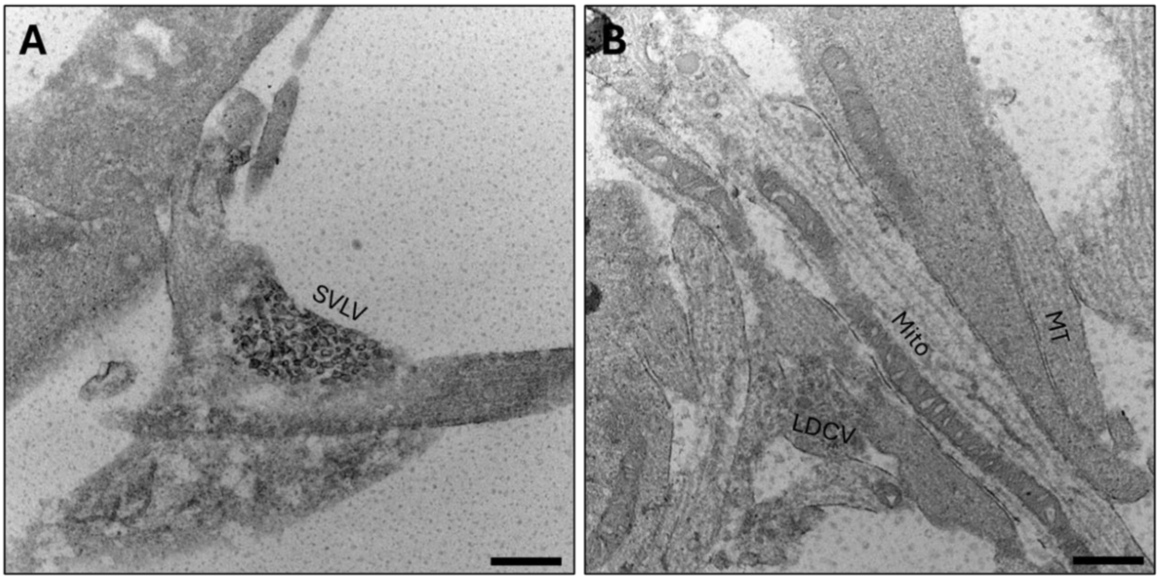


*Figure S6: Exemplary electron micrographs of SH-SY5Y cells differentiated with RA + BDNF and additionally Cholesterol A: Irregularly shaped synaptic-like vesicles in neurite extension lacking the characteristic synaptic structure of a postsynaptic counterpart. B: Axons with microtubules containing long tubular mitochondria, and LDCVs. Abbreviations: Mito: Mitochondria; MT: Microtubule; LDCV: large dense core vesicles; SVLV: Synapse-like synaptic vesicles. Scale bars: 500 nm.*

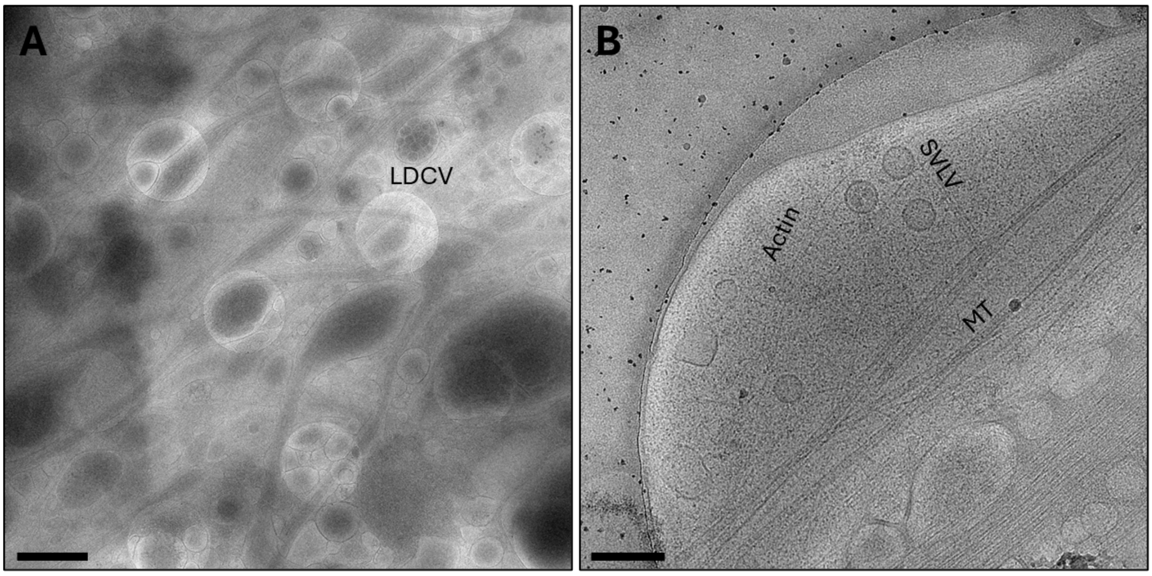


*Figure S7: Cryo-EM Micrographs of neurite extensions in SH-SY5Y cells at DPD28. A: Densely packed neurite network of SH-SY5Y cells. LDCVs can be seen in neurites. B: Varicosity in a neurite bulging out to the edges of a holey carbon grid. Scale bars: 2 µm (A), 100 nm (B)*

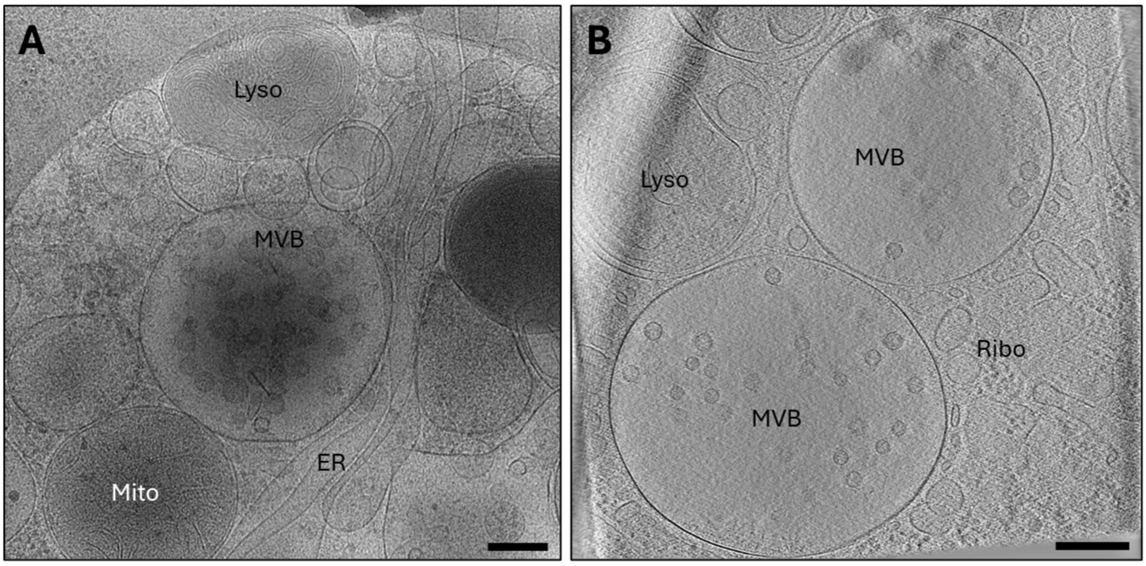


*Figure S8: Cryo-EM Micrograph (left) and Cryo-ET (right) of multivesicular bodies (MVB) containing protein decorated vesicles, which - along with the lack of connectors and an active zone - clearly differentiates it from a presynaptic bouton. Abbreviations: Mito: Mitochondria; ER: endoplasmatic reticulum; Lyso: lysosome; Ribo: Ribosome. Scale bar: 200 nm.*

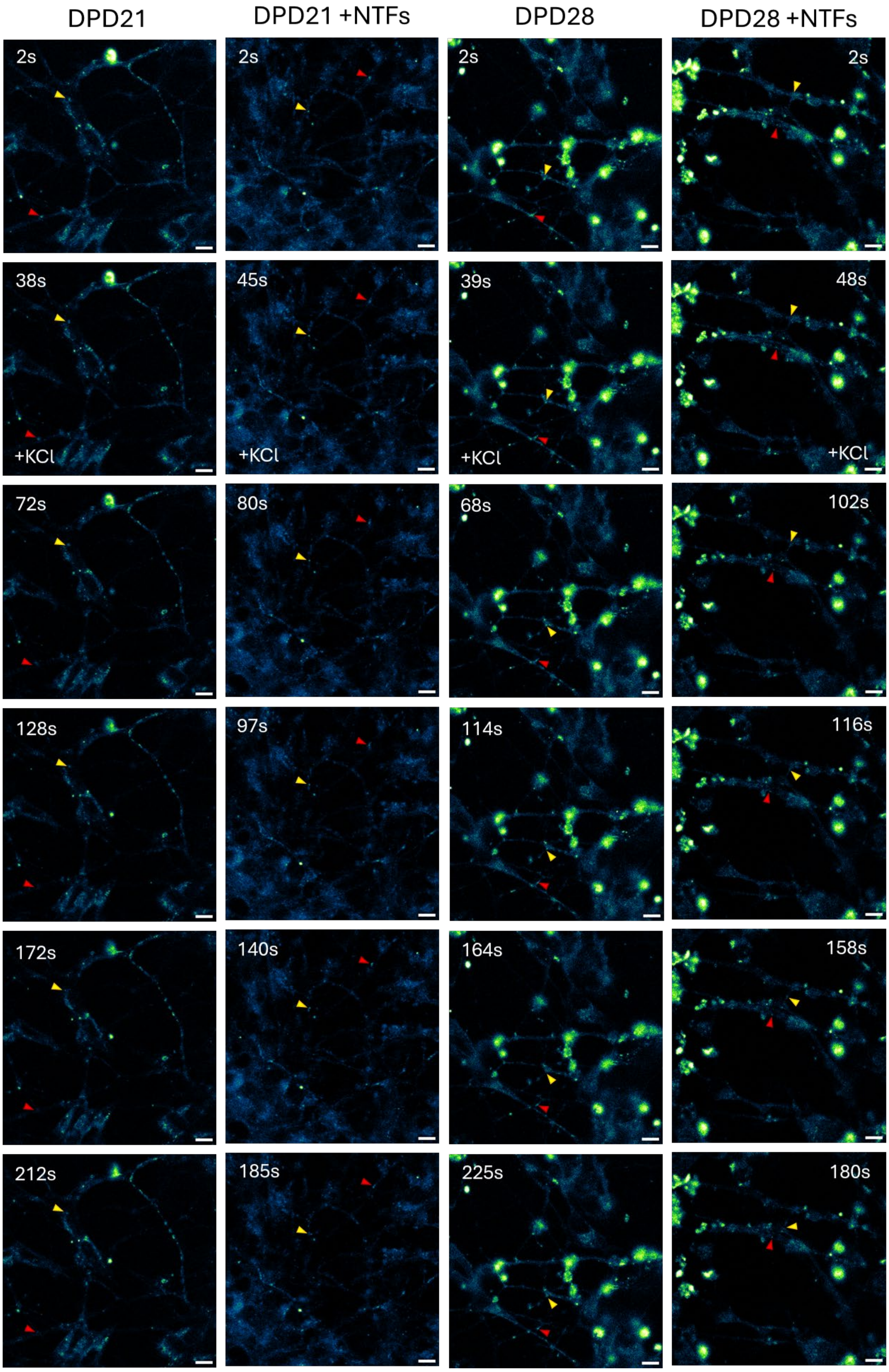


*Figure S9: Movement of fluorescent signal after AM4-64 staining. Selected time frames from the live staining experiment shown in Figure 5. SH-SY5Y cells at DPD 21 and DPD 28 were imaged under standard culture conditions or with additional NTFs. Fluorescence intensity is shown using the same pseudocolor scale as in Figure 5. Red and yellow arrows indicate two selected puncta tracked over time. The second frame was acquired directly after the addition of high-potassium Tyrode's solution (+KCl) to stimulate exocytosis. Anterograde and retrograde movements of puncta without fluorescence loss, regardless of stimulation, suggest the absence of stimulus-evoked synaptic vesicle exocytosis. Puncta move out of focal plane (red arrow in DPD21), but do not undergo exocytosis. Corresponding live movies in Figure S16-19. Scale bars: 10 µm.*

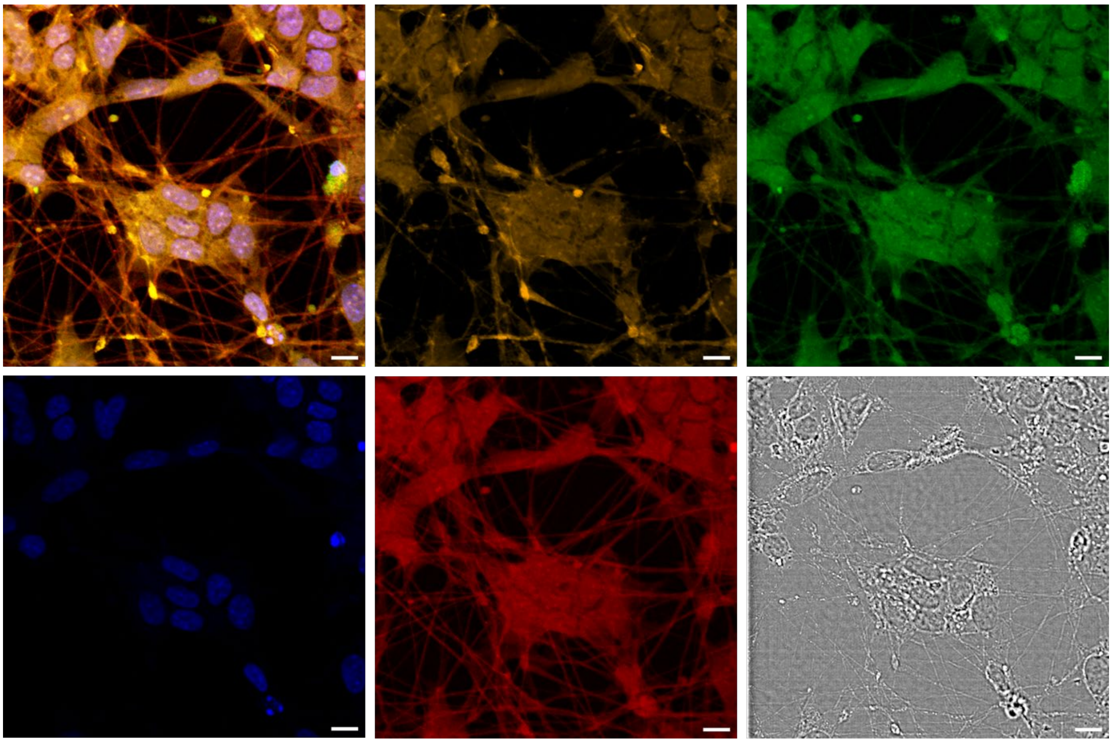

*Figure S10: Confocal image of SH-SY5Y cells at DPD 28 differentiated with additional NTFs. Synaptophysin (orange), PSD95 (green), DAPI (blue), β-tubulin III (red) and brightfield (grey). Scale bars: 20 µm.*

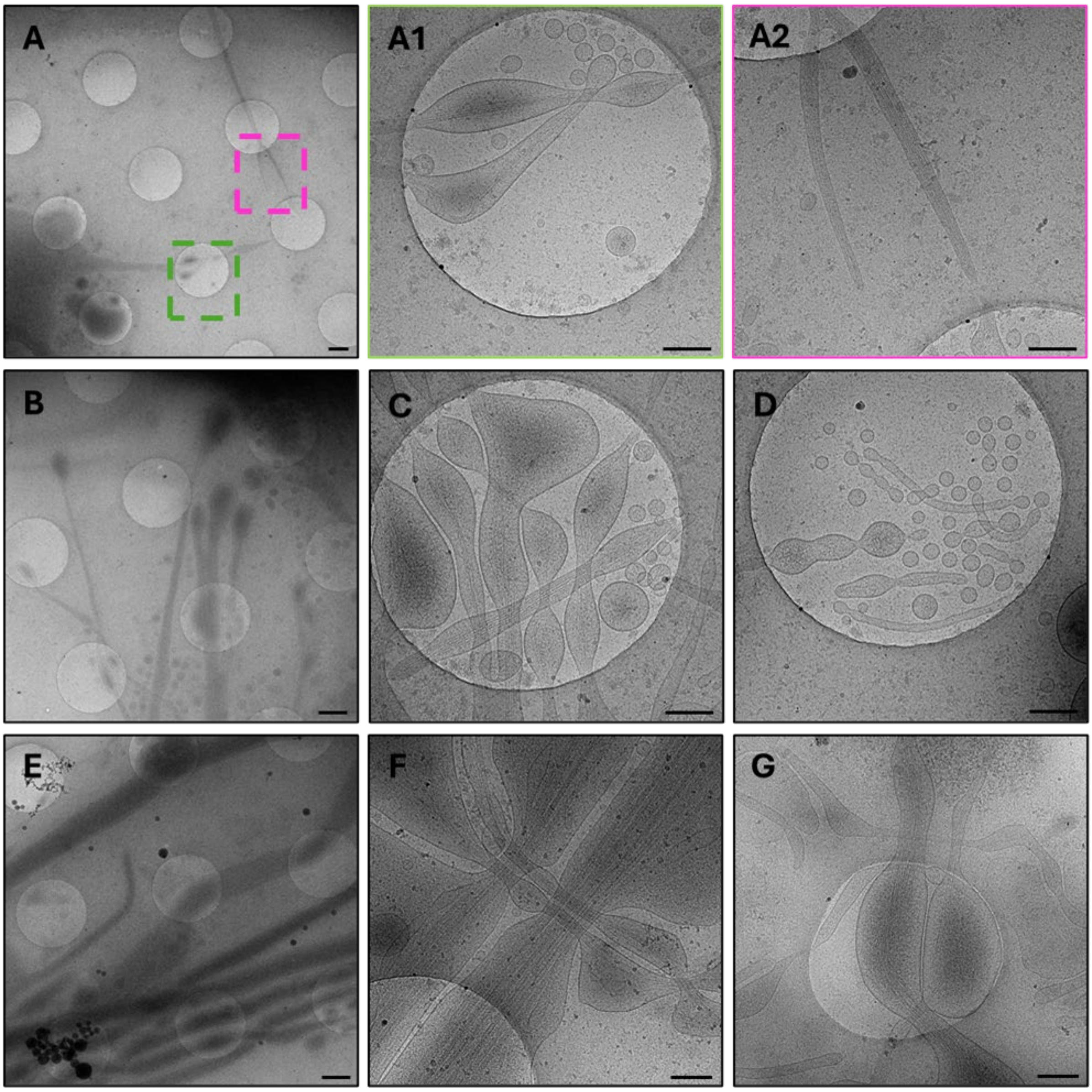


*S 11: Cryo-EM micrographs of thin projections in SH-SY5Y cells. Thin projections containing actin bundles are more frequent in early differentiation points and appear to be replaced with neurites containing microtubules upon further differentiation. A: SH-SY5Y cells at DPD 14 showing only few neurites that contain straight organized actin bundles characteristic of TNTs; A1: Zoom in of a possible TNT with actin structure and extracellular vesicles budding. A2: Zoom in of a branching extension with actin structure. B-G: thin TNT-like extensions and neurites at DPD 28. B: Overview of thin neurites of SH-SY5Y cells. C: TNT-like structures with bulging phenotype crossing each other. D: EV budding in TNTs-like extensions. E: Overview of TNT-like extensions and neurites in proximity. F: Axonal neurites with microtubules crossing thin TNT-like extensions. G: Thin TNT-like extensions bulging in holey carbon grids.*

Movies

S12: Movie of the tomogram seen in Figure 3A

S13: Movie of the tomogram seen in Figure 3B

S14: Movie of the tomogram seen in Figure 3C

S15: Movie of the tomogram seen in Figure 3D

S16-19: Movies of live-imaging seen in Figure 5 and S8

S20: Movie of the tomogram seen in Figure S8B
